# Idiopathic chronic diarrhea in rhesus macaques is not associated with enteric viral infections

**DOI:** 10.1101/2021.07.01.450785

**Authors:** Eric Delwart, Michael J. Tisza, Eda Altan, Yanpeng Li, Xutao Deng, J. Dennis Hartigan-O’Connor, Amir Ardeshir

## Abstract

While recent changes in treatment have reduced the lethality of idiopathic chronic diarrhea (ICD), this condition remains one of the most common causes of rhesus macaque deaths in non-human primate research centers. We compared the eukaryotic viromes in fecal swabs from 52 animals with ICD and 41 healthy animals. Viral metagenomics targeting virus-like particles was used to identify viruses shed by each animal. Five viruses belonging to the *Picornaviridae*, one to the *Caliciviridae*, one to the *Parvoviridae*, and one to the *Adenoviridae* families were identified. The fraction of reads matching each viral species was then used to estimate and compare viral loads in ICD cases versus healthy controls. None of the eukaryotic viruses detected in fecal swabs were strongly associated with ICD. Other potential causes of ICD are discussed.

## Introduction

Idiopapthic chronic diarrhea (ICD) is the leading cause of morbidity in captive colonies of rhesus macaques and a common cause of non-medical research mortality (1, 2) and is defined by chronic or recurring non-bloody diarrhea and microscopic colonic ulcers (2–5). ICD cases are typically diagnosed clinically before one year of age. The histopathological evaluations of the ICD cases show surface epithelium attenuation, goblet cell depletion and crypt branching and microscopic ulcers (6). The lamina propria is also filled with infrates including lymphocytes and plasma cells (5, 6). In addition, Rhesus macaques’ enterocolitis is a T_H_1 biased immune response with concurrent gut microbial dysbiosis (7–10).

Bacterial infections with *Shigella, Campylobacter, Yersinia, Salmonella, and/or Clostridium difficile* as well as parasitic infections, including *Cryptosporidium*, are common known causes of diarrhea in macaque colonies (2, 11). Cases of persistent and recurrent diarrhea in which no pathogenic bacteria, parasites or other etiological agents are found are defined as ICD (8).

Rhesus macaques with ICD respond poorly to treatment, including corticosteroids or antibiotics, resulting in frequent hospitalization from dehydration and weight loss and non-responsive animals are euthanized for ethical considerations (6).

A study comparing viral etiology of diarrheal diseases in rhesus macaques found that rhesus and pigtail macaques housed in the monkey farm of the Institute of Laboratory Animal Sciences, Chinese Academy of Medical Sciences shed enteroviruses, enteric adenoviruses, coronaviruses, and rotaviruses, while others at the Yerkes National Primate Research Center (Yerkes, GA) contained mainly enteroviruses and eneteric adenoviruses (12). Adenoviruses and rotaviruses were also associated with diarrhea in captive rhesus macaques (12–14). Inoculation with primate caliciviruses induced colitis in juvenile rhesus macaques (15). SIV-induced enteropathy was associated with a greater diversity of enteric viruses and parvovirus viremia when compared to SIV-negative rhesus macaques (16).

We previously described the virome in feces of healthy macaques as well as in those acute diarrhea or ICD and reported that shedding of several picornavirus genotypes were weakly associated with ICD while parvoviruses were weakly associated with healthy animals (17). Here we extend these studies by analyzing fecal samples from 52 animals with ICD and 41 healthy controls.

## Materials and methods

### Study design

This study is a case-control to search for the possible differences in the stool virome of ICD and healthy animals. The study was not blinded.

### Ethics

All experimental animal procedures were approved by the UC Davis Institutional Animal Care and Use Committee (IACUC Protocol #: 21763) and were in accordance with the requirements of the USDA Animal Welfare Act and the Guide for the Care and Use of Laboratory Animals and the research adhered to the Principals for the Ethical Treatment of Nonhuman Primates. Assurance Number D16-00272(A3433-01). All sample collection activities were conducted per standard operating procedures at the California National Primate Research Center (CNPRC). The UC Davis animal care program is fully accredited by the Association for the Assessment and Accreditation of Laboratory Animal Care, International (AAALACi), registered with USDA, and maintains a Public Health Services Assurance.

### Animals

All animals in this study were born, and raised at the CNPRC and samples collected while they are housed in oudoor colony (healthy controls) or from necropsy (ICD). The healthy control animals (n=41) were selected from outdoor housed animals without any gastrointestinal issues (including diarrheal diseases) or antibiotic treatment within the last 6 months prior to sampling. ICD cases (n=52) were selected from the cases that were hospitalized for the third time for non-pathogneic diarhea within the past 365 days and then euthanized. ICD cases that were euthanized with other accompanying diseases were excluded from the study to control for confounding factors.

### Sample collection

Rectal swabs were collection from rhesus macaques while sedated for the bi-annual physical exam (controls) or during necropsy (ICDs) and immediately frozen at −70°C.

### Generation of mNGS libraries

One mL of PBS buffer was added to fecal swabs, vigorously vortexed, and removed from the collection tube. The suspension (500 µl) was then placed in a microcentrifuge tube with 100 µl of zirconia beads and vigorously vortexed. The fecal suspension was centrifuged at 8000× *g* for 5 min at 4°C, and the supernatant (500 µL) was filtered through a 0.45 µm spin column filter (Millipore, Burlington, MA, USA) to remove bacteria and other large particulates. The filtrate was digested for 1.5 hours at 37°C with a mixture of nuclease enzymes consisting of 14U of Turbo DNase (Ambion, Life Technologies, USA), 3U of Baseline-ZERO (Epicentre, USA), 30U of Benzonase (Novagen, Germany), and 30U of RNase One (Promega, USA) in 1x DNase buffer (Ambion, Life Technologies, USA). Then 150 ul of the above mixture was extracted using the MagMAX™ Viral RNA Isolation kit (Applied Biosystems, Life Technologies, USA), and nucleic acids were resuspended in 50 µl water with 1 ul RiboLock RNAse inhibitor. 11 µl of nucleic acids were incubated for 2 min at 72°C with 100 pmol of primer A (5’GTTTCCCACTGGANNNNNNNN3’) followed by a reverse transcription step using 200 units of Superscript III (Invitrogen) at 50°C for 60 min with a subsequent Klenow DNA polymerase step using 5 units (New England Biolabs) at 37°C for 60 min. cDNA was then amplified by a PCR step with 35 cycles using AmpliTaq Gold™ DNA polymerase LD with primer A-short (5’GTTTCCCACTGGATA3’) at an annealing temperature of 59°C. The random amplified products were quantified by Quant-iT™ DNA HS Assay Kit (Invitrogen, USA) using Qubit fluorometer (Invitrogen, USA) and diluted to 1 ng of DNA for library input. The library was generated using the transposon-based Nextera™ XT Sample Preparation Kit using 15 cycles (Illumina, San Diego, CA, USA) and the concentration of DNA libraries was measured by Quant-iT™ DNA HS Assay Kit. The libraries were pooled at equal concentration and size selected for 300 bp – 1,000 bp using the Pippin Prep (Sage Science, Beverly, MA, USA). The library was quantified using KAPA library quantification kit for Illumina platform (Kapa Biosystems, USA) and a 10 pM concentration was loaded on the MiSeq sequencing platform for 2×250 cycles pair-end sequencing with dual barcoding.

### Bioinformatics

Human and bacterial reads were identified and removed by comparing the raw reads with human reference genome hg38, and bacterial genomes release 66 (collected from ftp://ftp.ncbi.nlm.nih.gov/blast/db/FASTA/, Oct. 20, 2017) using local search mode. The remaining reads were de-duplicated if base positions 5 to 55 were identical. One random copy of duplicates was kept. The sequences were then trimmed for quality and adaptor and primer sequences by using VecScreen. (18) After that, the reads were de novo assembled by Ensemble Assembler (19). Assembled contigs and all singlet reads were aligned to an in-house viral protein database (collected from ftp://ftp.ncbi.nih.gov/refseq/release/viral/, Oct. 20, 2017) using BLASTx (version 2.2.7) using E-value cutoff of 0.01. The significant hits to the virus were then aligned to an in-house non-virus-non-redundant (NVNR) universal proteome database using DIAMOND (10) to eliminate the false viral hits. Hits with a more significant (lower) E-value to NVNR than to the viral database were removed. The remaining singlets and contigs were compared to all eukaryotic viral protein sequences in GenBank’s non-redundant database using BLASTx [23]. The genome coverage of the target viruses was further analyzed by Geneious R11.1.4 software (Biomatters, New Zealand Quantitation of viral reads: Genetic variation was observed between homologous sequences belonging to the same enterovirus genotypes. For estimating the viral abundance of each viral types using the number of viral reads, closely related contigs (of the same viral genotype) from multiple animals were concatenated (with 100 bp ‘N’ spacers to prevent reads aligning to the areas where the contigs are joined)(Supplemental Table 1 and Dataset 1). These concatenated contigs where then used to find matching reads using the nucleotide aligner program Bowtie2 with the seed length parameter set at 30, and the alignment was considered a hit if the read identity to the reference concatamer was greater than 95%. A total of 11 concatenated contigs representing the most commonly detected viruses were generated. For normalization (to account for the variable number of total reads generated from different samples) the number of matching read hits were converted to reads per million total reads (RPM). Heat map was generated using freeware program at heatmapper.ca and the expression function. The RPM were used as input. For clustering method the default average linkage was used. For distance measurements Euclidian distances were used. Both row and column dendograms are shown. The z-score is the number of standard deviations by which the value of a raw score is above or below the mean value of viral reads per millions (RPM).

### Quantitation of de novo assembled virus contigs

The protocols used to predict virus contigs with Cenote-Taker 2 have been described (20, 21) by running Cenote-Taker 2 (version 2.0.0) with a minimum contig size of 500 nucleotides and the “standard” database. Contigs were dereplicated using the Mummer-based RedRed program (22) (https://github.com/kseniaarkhipova/RedRed). Low-complexity regions of contigs were masked with RepeatMasker (23). Abundance was calculated with the same methods as described (21). Finally, sample-by-sample swarm plots were generated using DaBest python package (24).

## Results

Fecal swabs were collected from 52 macaques suffering from ICD and 41 healthy controls. Fecal material was filtered to enrich for virus-like particles. Host and bacterial nucleic acids were then depleted using nuclease enzymes. Total nucleic acids were extracted from digested and filtered materials and amplified using random RT-PCR. DNA was then converted to Illumina-compatible DNA and sequenced (see materials and methods). All raw data are available in GenBank under Bioproject PRJNA608547. The resulting reads were analyzed using BLASTx to identify reads matching all known eukaryotic viruses. Eight different simian viruses were identified. The most prevalent viruses belonged to the *Picornaviridae* family, including members of enterovirus species A (enterovirus 19, 46, 92), enterovirus species J (enterovirus 103), and sapelovirus (species B). Also identified but found in fewer animals were sapovirus (*Caliciviridae*), adenovirus (*Adenoviridae*), and bocavirus (*Parvoviridae*). The percent of reads matching these different simian viruses was then plotted in order to compare the eukaryotic virome composition of ICD cases to that of healthy animals (Fig 1).

**Figure 1.**
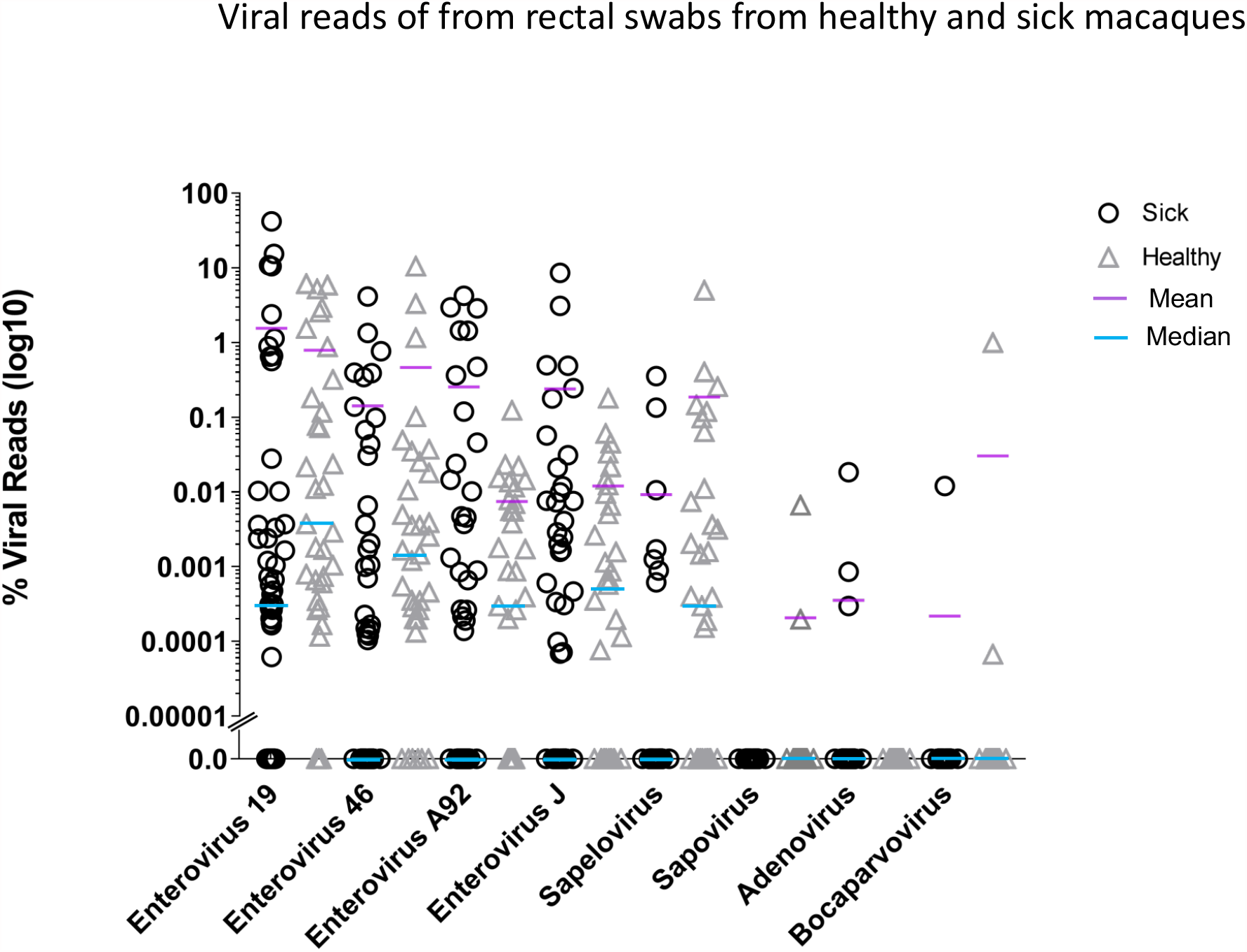
Fraction of viral reads matching different viruses are compared between fecal swabs from ICD cases versus healthy animals. Read numbers were quantified as described in materials and methods.

This analysis indicated that for the viruses detected in most animals (five picornaviruses), the median and mean read percentages were higher in healthy than sick animals, with the single exception of enterovirus A92 whose mean value in ICD animals was above that of the healthy control group.

When the data for these five most common infections was plotted using whisker plots, no difference between the ICD and healthy cases could be seen, except again for mean enterovirus A92 representation being lower in healthy animals (Fig 2).

**Fig 2.**
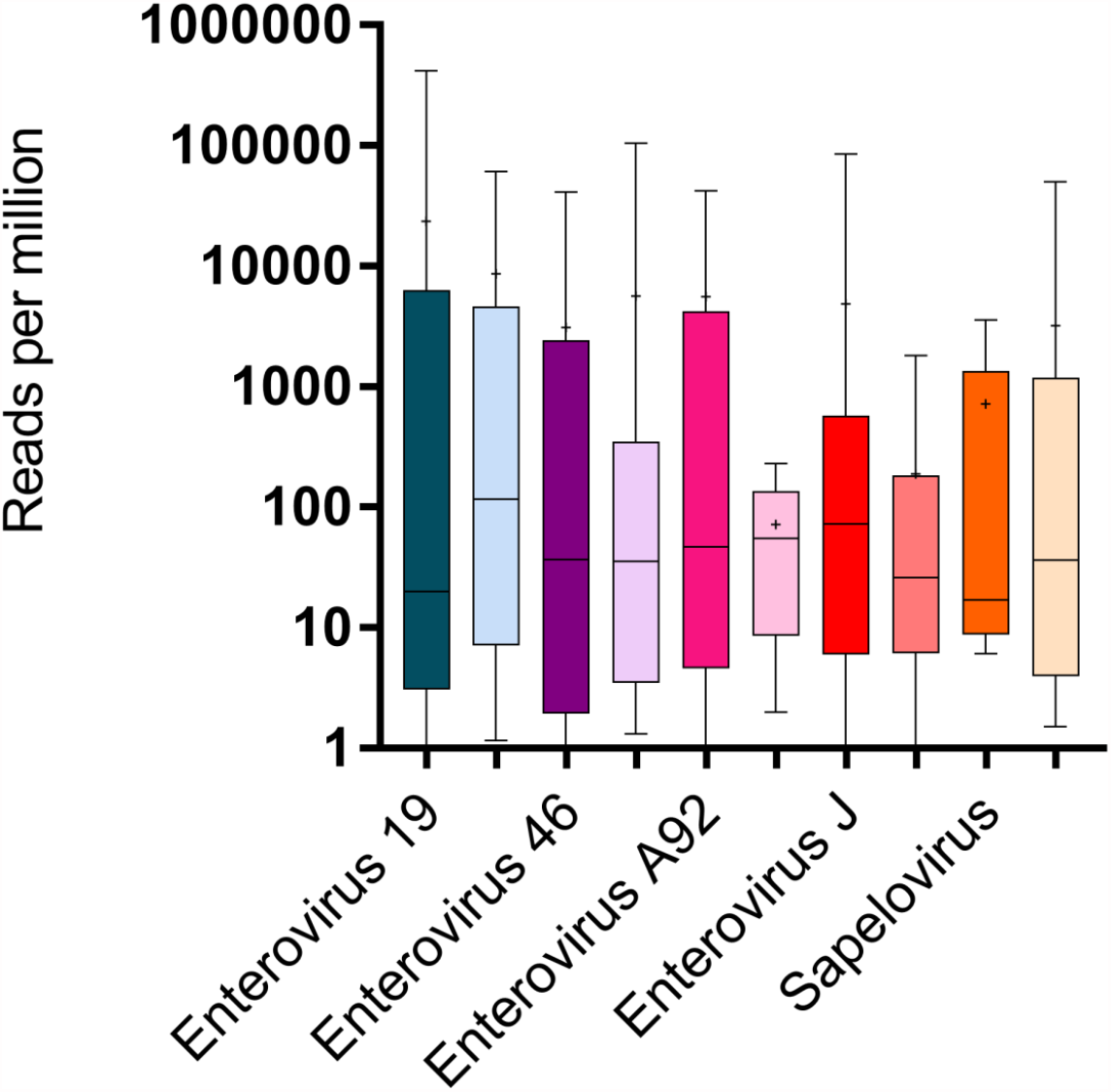
Whisker plot of the RPM for the most common viruses. Lighter color bargraphs are from healthy control animals.

**Fig 3.**
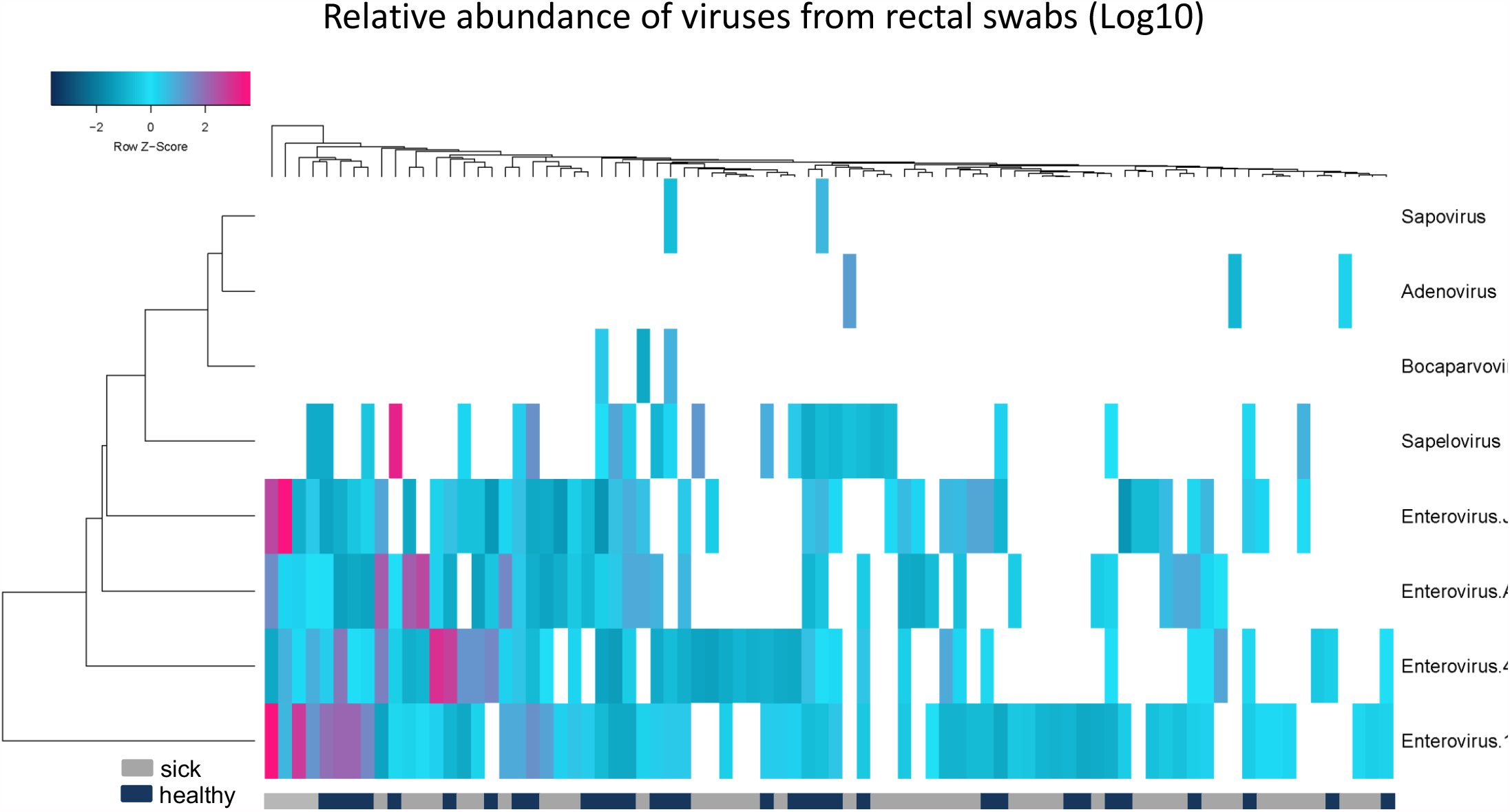
Unsupervised hierarchical clustering analysis of fecal swab viromes suggests that there are no distinct divisions between the cases and control subjects.

Next a hierarchical clustering program (heatmapper.ca) was used to determine if the viromes of animals with or without ICD clustered separately. No clustering of related viromes based on clinical status (Gray and black bar below heat map) was detected further supporting the lack of association between ICD and virome composition.

We next used Cenote-Taker 2 to identify viral contigs from the Illumina libraries from each animal in the two macaque groups. Cenote-Taker 2 scans open reading frames of contigs for virus hallmark genes, identifying known and divergent viral contigs using a curated dataset and enabling the discovery and annotation of both DNA and RNA viral genomes (20, 21). Such hallmark genes include structural genes such as capsid proteins, genome-packaging genes such as terminases, and genome-replication genes such as RNA-dependent RNA polymerase and replicative helicase (20, 21). The abundance of these viral contigs was compared between ICD and healthy animals by measuring reads per kilobase per millions reads (RPKM) (Figure 4). Only contigs that met minimum thresholds of 0.05 reads per kilobase (RPKM) of transcript in at least 10% all samples were compared in this analysis, as rare contigs are unlikely to originate from viruses with a frequent role in ICD pathogenesis.

**Figure 4:**
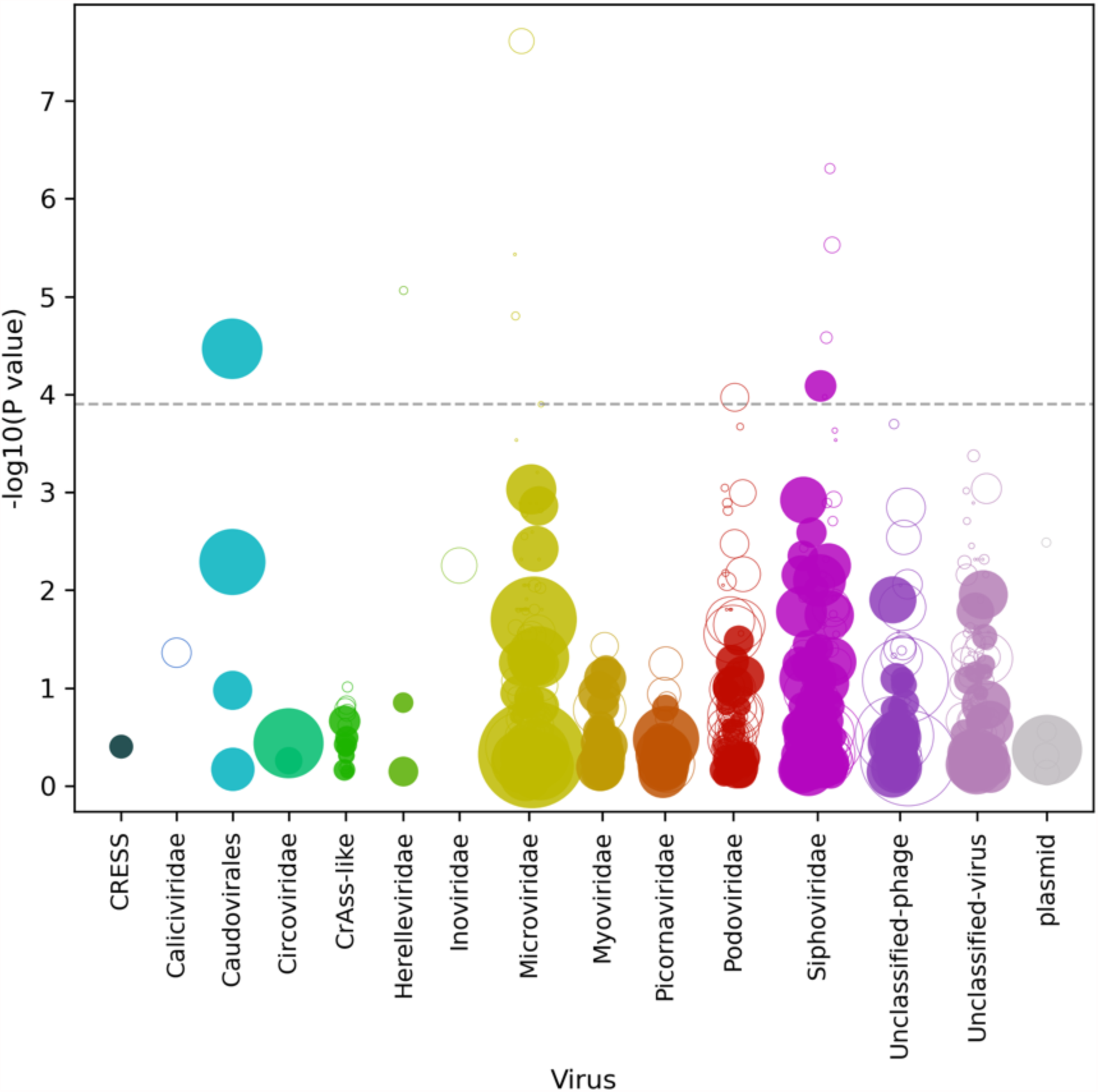
Virome-wide associations of macaques with ICD represented as a Manhattan plot. Each virus contig is represented as a dot along the x-axis, with its y-axis value being the inverse log10 P value. The size of each dot corresponds to the median relative abundance of the taxon in the disease cohort. Filled dots represent virus contigs found at higher abundance in macaques with ICD while hollow dots represent decreased abundance in macaques with ICD. The dashed gray line represents a false discovery rate threshold of 1%, calculated using the Benjamini-Hochberg procedure.

The 20 virus contigs with the lowest p values (i.e. highest −log_10_p value) were then compared at a sample-by-sample level (Fig 5A). While twelve virus contigs were significantly different between cohorts after multiple testing correction, it was observed that most relationships represented a minority of samples from one of the cohorts being present at very high RPKM values. Finally, sample-by-sample comparison of eukaryotic viruses (threshold of 10% of samples > 0.05 RPKM) showed no population level differences (Fig. 5B). Comparison of abundance of de novo assembled contigs is therefore consistent with the other analyses showing that none of the eukaryotic viruses are associated with ICD.

**Figure 5:**
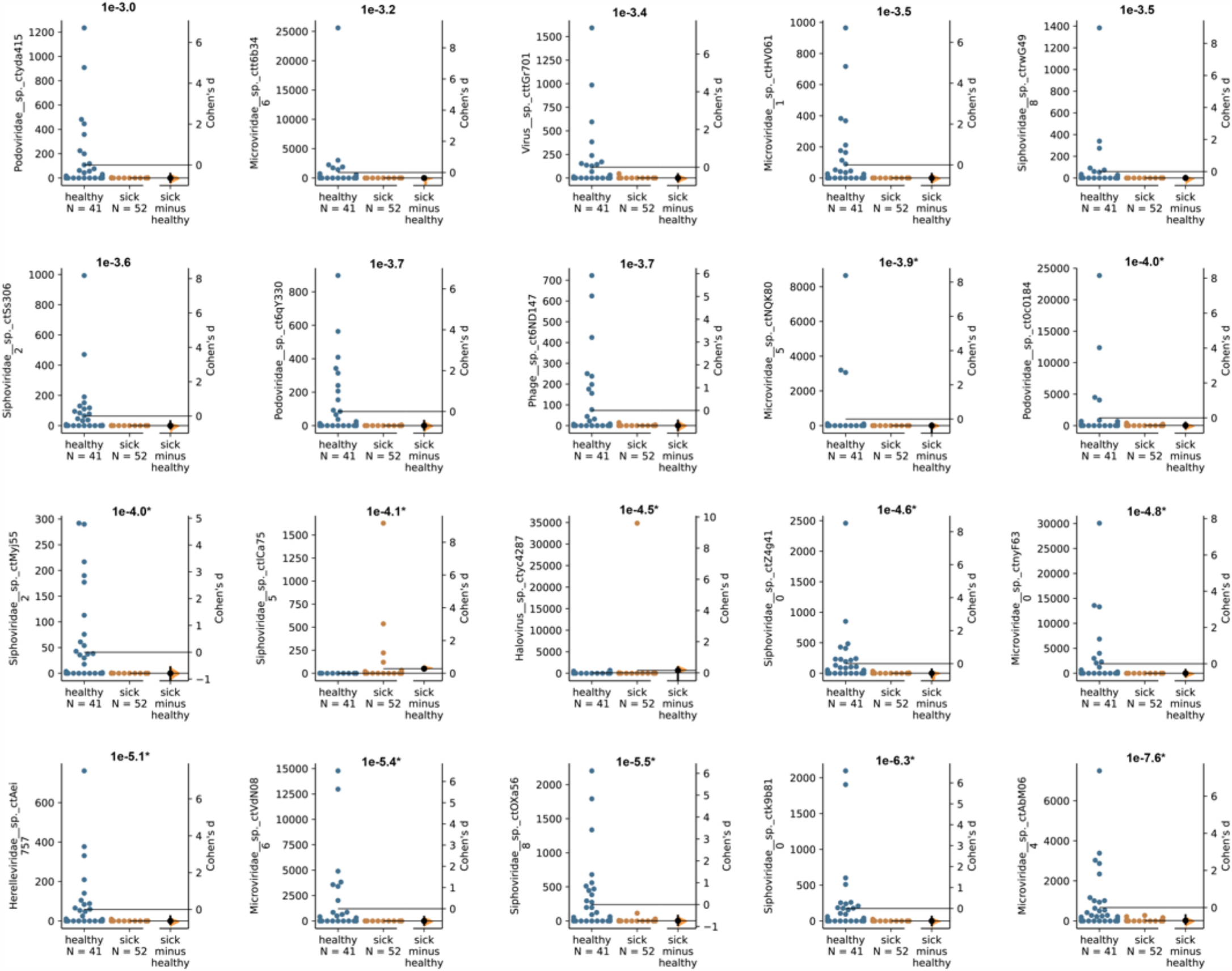
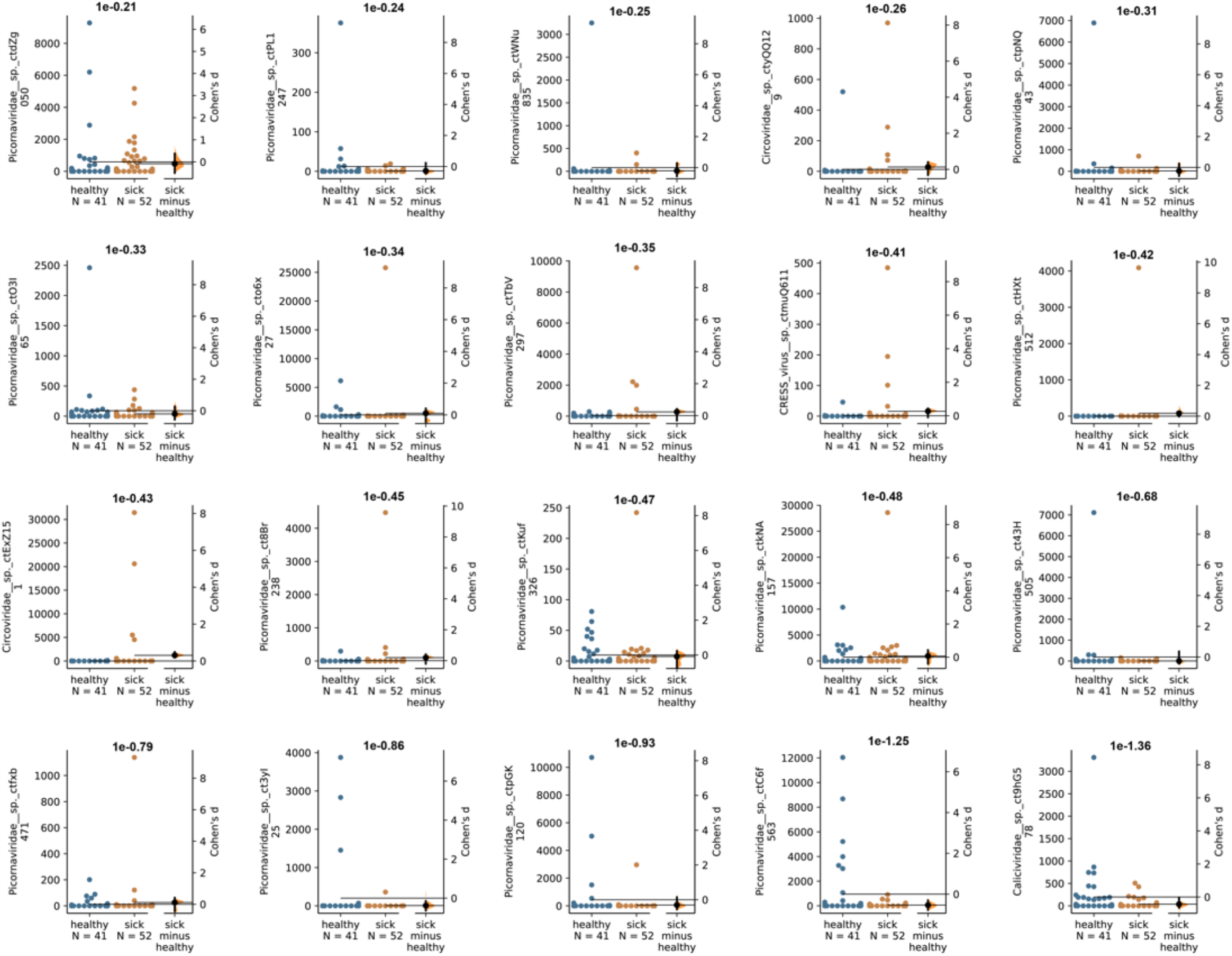
Sample-by-sample comparison of viruses with largest differences between healthy and ICD cohorts. Each plot represents a different virus contig, with RKPM values from healthy (blue) and ICD (orange) samples displayed. Cohen’s d effect size values are plotted on the right side of each plot, and uncorrected p values are reported above each plot. Asterisks (*) next to −log_10_p value on top of each graph were given to virus contigs with significant p values after multiple testing correction. (A) 20 virus contigs with the lowest p value overall. (B) Eukaryotic virus contigs with lowest p value. Note that most picornaviruses contigs retrieved from de novo assembly represented sub-genomic fragments with >90% average nucleotide identity to previously discovered macaque viruses.

## Discussion

We tested the hypothesis that a specific eukaryotic virus was associated with ICD. We used sequence homology of NGS reads derived from viral metagenomics to identify viruses in fecal swabs. The frequencies of reads to different viruses were then used as surrogates for viral loads. By comparing the viruses in rhesus macaques with ICD versus those in healthy controls, we were unable to associate any eukaryotic virus with ICD. These results differ from the conclusions of a previous analysis, in which we showed that some enterovirus genotypes were weakly associated with ICD (17). During the interval between these studies the treatment for ICD changed and the frequency of ICD dropped significantly. Specifically, antibiotic therapy was no longer done on animals testing negative for enteropathogens; these animals instead received fluid therapy, and synbiotic or probiotic treatments. The ICD fecal swabs samples that were analyzed in this study were collected during necropsy from severe cases that required medical cull while feces analyzed in prior studies were feces collected from cage pans while the animals were still under treatment. Therefore, it is possible that during the later stage of ICD studied here, shedding of enteric viruses in watery diarrhea is reduced due to more extensive damage to the lining of the gut and consequent reduction in viral target cells. It is possible that earlier sampling of ICD-associated enteric virome may be necessary to identify viruses associated with this condition.

We observed that the median level of viral shedding was generally higher in healthy than in ICD cases (Fig 1 and 2), a result consistent with a reduction in viral target cells in diarrheic animals. In the analysis of virus contigs predicted by Cenote-Taker 2 (Figs. 4 and 5), it was also observed that fewer reads to specific phages were generally found in swabs from ICD cases versus healthy animals. Such a reduction in phage shedding may reflect a gut content with fewer bacterial hosts as its content is flushed out due to chronic diarrhea.

## Acknowledgments

We thank E. Fahsbender for technical help and data generation and R. Bruhn for assistance for statistical analyses. We thank CNPRC for assistance with sample collection.

